# The paternally imprinted gene *Snord116* regulates cortical neuronal activity

**DOI:** 10.1101/809822

**Authors:** Pace Marta, Colombi Ilaria, Falappa Matteo, Andrea Freschi, Mojtaba Bandarabadi, Armirotti Andrea, Blanco María Encarnación, Antoine R. Adamantidis, Amici Roberto, Cerri Matteo, Chiappalone Michela, Tucci Valter

## Abstract

Prader-Willi syndrome (PWS) is a neurodevelopmental disorder that is characterized by rapid eye movement (REM) sleep abnormalities. The disease is caused by genomic imprinting defects that are inherited through the paternal line. Among the genes located in the PWS region on chromosome 15 (15q11-q13), small nucleolar RNA 116 (*Snord116*) has been previously associated with intrusions of REM sleep into wakefulness in both humans and mice.

Here, we further explore the processes of sleep regulation by studying the PWScr^m+/p-^ mouse line, which carries a paternal deletion of *Snord116.* We focused on microstructural electrophysiological components of sleep, such as REM sleep features and sleep spindles within NREM sleep. While the former are thought to contribute to neuronal network formation early in brain development, the latter are markers of thalamocortical processes. Both signals are often compromised in neurodevelopmental disorders and influence functional properties of cortical neurons. Thus, we isolated and characterized the intrinsic activity of cortical neurons using *in vitro* microelectrode array (MEA) studies.

Our results indicate that the *Snord116* gene in mice selectively influences REM sleep properties, such as theta rhythms and the organization of REM episodes throughout sleep-wake cycles. Moreover, sleep spindles present specific abnormalities in PWS model systems, indicating that these features of sleep may translate as potential biomarkers in human PWS. We observed abnormalities in the synchronization of cortical neuronal activity that are accounted for by high levels of norepinephrine.

In conclusion, our results provide support for an important role of *Snord116* in regulating brain activity during sleep and, in particular, cortical neuronal properties, thereby opening new avenues for developing interventions in PWS.

**Significance Statement:** We found that the *Snord116* gene, a major player in Prader-Willi syndrome (PWS), significantly impacts REM sleep and its regulation. Additionally, we found that sleep spindles, a subtle electroencephalography (EEG) marker that occurs during NREM sleep, are dysregulated in PWS mice that carry a paternal deletion of the *Snord116* gene. Using a combination of *in vivo* and *in vitro* experiments, we identified sleep features at the network and molecular level that suggest that *Snord116* is fundamental in the synchronization of neuronal networks. Our study also provides a new pre-clinical tool to investigate the pathophysiology of sleep in PWS.

## Introduction

Sleep and circadian alterations are frequently reported in subjects with Prader-Willi syndrome (PWS), representing a burden that impacts the quality of life for patients and their families. Indeed, up to 76% of PWS patients exhibit abnormal sleep (Gunay-Aygun, Schwartz, Heeger, O’Riordan, & Cassidy, 2001; Williams, Scheimann, Sutton, Hayslett, & Glaze, 2008). PWS is a neurodevelopmental disorder characterized by hypotonia and failure to thrive in infancy. From approximately 2 years of age, the phenotype shifts to hyperphagia, which leads to obesity (Holland, Treasure, Coskeran, & Dallow, 1995; Tauber, Thuilleaux, & Bieth, 2015). Other phenotypic hallmarks of PWS include mild to moderate cognitive deficits, behavioural problems including obsessive–compulsive disorder, and sleep disturbances. Among these sleep disturbances, PWS subjects exhibit excessive daytime sleepiness (EDS) (Camfferman, McEvoy, O’Donoghue, & Lushington, 2008; Helbing-Zwanenburg, Kamphuisen, & Mourtazaev, 1993) and rapid eye movement (REM) sleep dysregulation (Bruni, Verrillo, Novelli, & Ferri, 2010; Vela-Bueno et al., 1984; Vgontzas et al., 1996). Such REM dysregulation implies a disruption in the circadian rhythms, producing an irregular sleep-wake cycle, and may be linked to the abnormal thermoregulation observed in PWS subjects (Clarke, Waters, & Corbett, 1989). Some studies have also reported narcoleptic-like symptoms such as sleep attacks, cataplexy and a transient loss of muscle tone (Tobias, Tolmie, & Stephenson, 2002; Vgontzas et al., 1996).

PWS is caused by the lack of expression of paternally imprinted genes in the chromosomal region 15q11-q13 The small nucleolar RNA (snoRNA) clusters in this region have a pivotal role in the aetiology of the disease. Individuals lacking the SNORD116 snoRNA cluster and the IPW (imprinted In Prader-Willi Syndrome) gene suffer the same failure to thrive, hypotonia, and hyperphagia that are observed in subjects with larger deletions and maternal uniparental disomy (de Smith et al., 2009; Sahoo et al., 2008). Additionally, it has been shown that mice with a paternal deletion of the *Snord116* gene (PWScr^m+/p-^) recapitulate the major human endophenotypes, including sleep alteration coupled with dysregulation of diurnal clock genes (Lassi et al., 2016; Powell & LaSalle, 2015).

We have previously described sleep alterations in PWScr^m+/p-^ mice, mostly related to REM sleep (Lassi et al., 2016). Here, we explored a specific feature of sleep microstructure concerning the processes that regulate REM sleep in PWScr^m+/p-^ mice. Next, we investigated sleep spindles, hallmarks of NREM sleep that have traditionally been challenging to characterize in mouse studies. Spindle activity is attributed to interactions between thalamic reticular, thalamocortical, and cortical pyramidal networks (Steriade, McCormick, & Sejnowski, 1993). Dysregulation of spindle properties is a sensitive indicator of thalamocortical and neuromodulator dysfunction in many diseases, including neurodevelopmental disorders (Krol, Wimmer, Halassa, & Feng, 2018). Previous studies have reported a complete absence or dysregulation of sleep spindles in PWS subjects (Gilboa & Gross-Tsur, 2013; Hertz, Cataletto, Feinsilver, & Angulo, 1993). Since children with PWS have been shown to have abnormal functional connectivity among brain regions (Zhang et al., 2013), a phenomenon that can derive from cortical and/or subcortical structures, these alterations may be responsible for the sleep alterations observed in PWS. Therefore, we sought to investigate the regulation of sleep by analysing electrophysiological elements dependent on the thalamocortical projections and intrinsic cortical processes. To investigate the properties of intrinsic cortical connectivity, with no interference from external stimuli and from the thalamocortical projections (Hinard et al., 2012), we studied *in vitro* cultures of dissociated cortical neurons using microelectrode array (MEA) recordings.

Our results confirm the function of the *Snord116* gene in sleep regulation and extend our understanding of sleep in PWS, for the first time, to the microstructure of sleep, a fine regulatory cortical process that controls sleep architecture.

## Materials and Methods

### In vivo study

#### Animals

Animal procedures were performed at the Istituto Italiano di Tecnologia (IIT) Genova and were approved by the Animal Research Committee and the Veterinary Office of Italy. All efforts were made to minimize the number of animals used, as well as any pain and discomfort, according to the principles of the 3 Rs (replacement, reduction, and refinement) (Tornqvist et al., 2014).

Mice carrying a paternal deletion of the *Snord116* gene and IPW exons A1/A2 B (Skryabin et al., 2007) (PWScr^m+/p-^) were used in this study, along with wild-type mice (PWScr^m+/p+^). In order to maintain the colony, patrilineal descendants of the original transgenic mice were bred on a C57BL/6J background. For the genotyping protocol, see the supplementary methods.

All animals used in this study were housed under controlled temperature conditions (22 ± 1 °C as room temperature) on a 12 h:12 h light/dark cycle (lights on 07:00–19:00). Food (standard chow diet) and water were provided ad libitum.

#### Experimental design

For the study, male mice with paternally inherited *Snord116* deletions (PWScr^m+/p-^) and wild-type littermate control mice (PWScr^m+/p+^) aged 15 to 18 weeks were used. To investigate the impact of *Snord116* in the sleep architecture, we made continuous recordings by electroencephalography combined with electromyography (EEG/EMG) for 24 h (12 h:12 h light/dark cycle, Figure 1A). Subsequently, we investigated changes in spindle properties at specific time points across the sleep-wake cycle. Specifically, EEG/EMG recordings were made for 2 h at two different time points as the baseline condition (B1 and B2; Figure 1D) to investigate spindle properties related to (i) circadian variation (switch from light to dark periods) and (ii) homeostatic variation (low and high sleep pressure). Thus, we investigated the frequency of sleep spindles and their properties in response to perturbation of sleep by performing 6 h of total sleep deprivation (SD). EEG/EMG recordings were made over the first hour of the rebound period (RB, Figure 1D) following 6 h of SD (RB1) and at two other time points over the 18 h recovery period (RB2 and RB3, Figure 1D).

**Figure 1.**
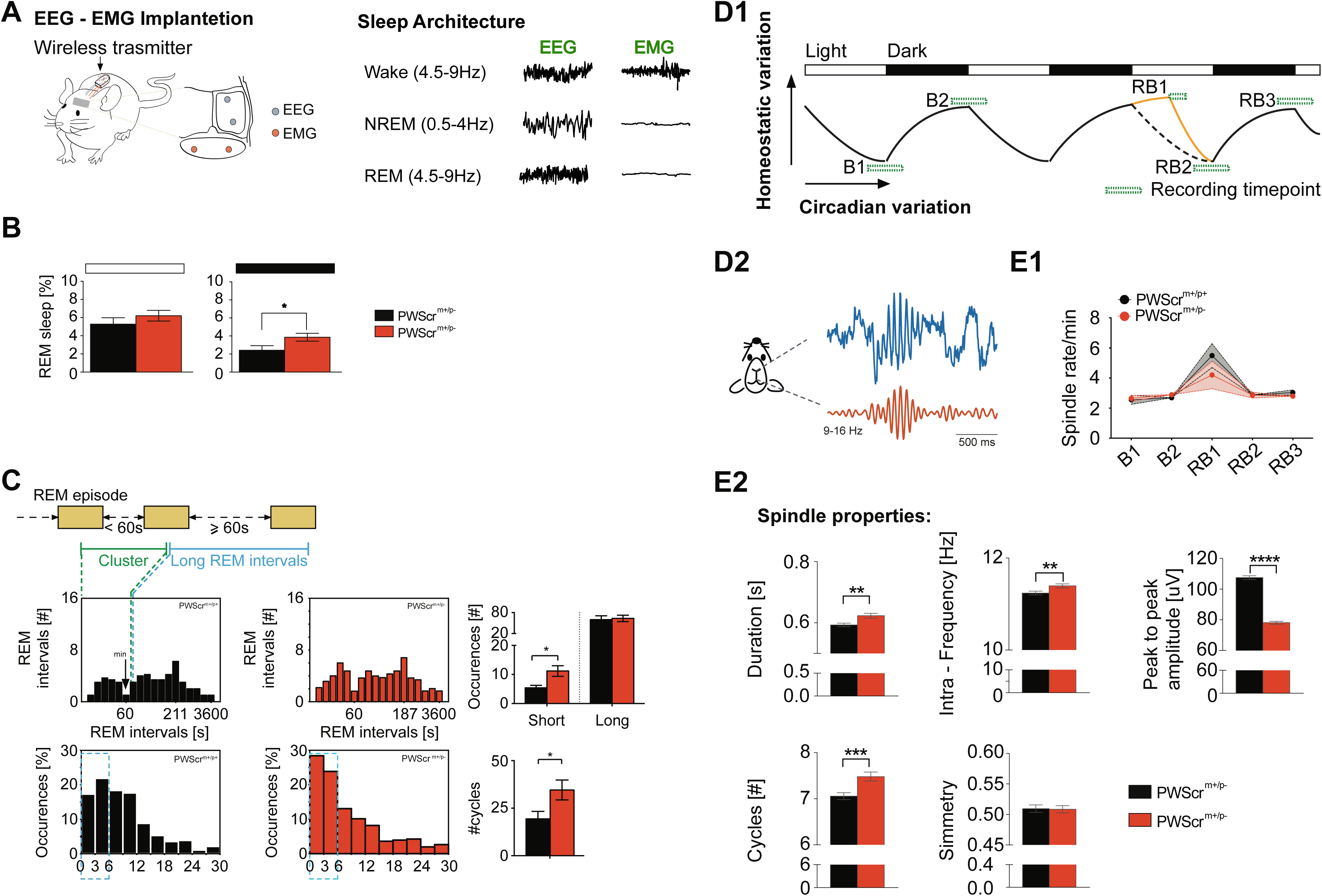
Paternal *Snord116* deletion impacts REM sleep and NREM sleep spindles: **A)** Schematic representation of EEG/EMG implantation to record the sleep-wake cycle and the recorded sleep architecture. With this system is possible to discriminate sleep-wake changes based on the frequency of EEG and the amplitude of the EMG signals. **B)** The percentage of time spent in in REM sleep during the 12 h light and dark period for PWScr^m+/p-^ mice (red) versus controls (black). Values are the 12 h means ± SEM. **C)** On the top, schematic representation of the criteria for the recognition of short-RSI and long-RSI. On the middle average frequency distribution of the duration of the interval from the end of one REM sleep episode to the beginning of the next REM sleep episode (REM sleep interval, RSI) recorded over the 24 of the sleep-wake cycles in PWScrm^+/p+^ (n= 10 in black) and PWScrm^+/p-^ mice (n= 10 in red). The frequency class which was taken as the boundary separating short and long-interval populations (minimum) is at 55-sec. Right, average value of the occurrence of the short and long-RSI for each genotype, values are expressed as mean ± SEM. On the bottom, histograms of sleep cycle length during the 24h of the sleep-wake cycle in PWScrm^+/p+^ (n= 10 in black) and PWScrm^+/p-^ mice (n= 10 in red). Sleep cycles are plotted for 3-hour bins. Only data up to 30 min are shown. Four combinations of criteria (see Method section). Cycles containing wake sequences >64 s were discarded from the analysis. On the right the average value of the number of the cycle length around 2-5 minutes in both genotypes of mice. **D1)** Schematic representation of experimental design to record sleep spindles properties. Sleep spindles were recorded at different time point according to the circadian and homeostatic sleep control. At the baseline sleep spindles were recorded at two time points (B1 and B2) for 2h between the shift from the light and dark period and vice versa, indicated by a green square. Next, sleep spindles were recorded over the first hour of the rebound period (RB1) following 6h of total-sleep deprivation (SD) and in other two-time points over the 18h of recovery period (RB2 and RB3). **D2)** Schematic representation of the spindles signal; **E1)** The number of the spindles estimated over the 24h of the sleep-wake cycles in both genotypes, PWScrm^+/p+^ (n= 10 in black) and PWScrm^+/p-^ mice (n= 10 in red). **E2)** Sleep spindles properties: duration between 0.4-2 s, central frequency, peak-to-peak amplitude, number of cycles between 5-30 and symmetry. Values are expressed as mean ± SEM. *p < 0.05; **p < 0.01; ***p < 0.001

#### *In vivo* SD protocol

Total SD was performed by gentle handling techniques, consisting of introducing novel objects into the cage or agitating (knocking or shaking) the cage when behavioural signs of sleep were observed. Animals were subjected to SD during the first 6 h of the light period, when mice usually sleep and are under high sleep pressure.

#### EEG/EMG electrode implantation

A total of 28 mice were used for the *in vivo* sleep study (n=14 PWScr^m+/p-^ and n=14 PWScr^m+/p+^ mice). Each mouse was anaesthetized using 1.5%–2.5% isoflurane in oxygen and surgically implanted with a telemetric transmitter (volume, 1.9 cm 3; total weight, 3.9 g; TL11M2-F20-EET; DSI, St. Paul, MN, USA) connected to electrodes for continuous EEG/EMG recording to assess the sleep-wake cycle. A wireless EEG transmitter/receiver was subcutaneously implanted. EEG wire electrodes were implanted epidurally over the right frontal cortex (coordinates: 2 mm posterior to bregma and 2 mm lateral to the midline, in the right frontal part of the skull) and the right parietal cortex (coordinates: 3 mm anterior to lambda and 2 mm lateral to the midline, in the right parietal part of the skull). EMG was recorded with 2 stainless steel wires inserted bilaterally into the neck muscles ∼5 mm apart and sutured in place (Figure 1A). Following surgery, all animals received paracetamol (200 mg/kg by mouth once a day; Tempra) and enrofloxacin (10 mg/kg subcutaneously once a day; Baytril) for two days after surgery. The animals were housed individually in their home cages for a recovery period of 7 days, after which EEG/EGM signals were recorded continuously for each mouse.

#### EEG/EMG analysis

Cortical EEG and EMG signals were recorded using Dataquest A.R.T. (Data Science International). Activity data measured only relative movement, which is dependent on the orientation and distance between the transmitter and receiver. Signals were digitized at a sampling rate of 500 Hz with a filter cut-off of 50 Hz. EEG signals were filtered at 0.3 Hz (low-pass filter) and 0.1 kHz (high-pass filter), respectively. The polysomnographic recordings were visually scored offline using SleepSign software (Kissei Comtec Co. Ltd, Japan) in four-second epochs to identify the wakefulness (W), NREM sleep (NR) or REM sleep (R) stages as previously described (Pace, Adamantidis, Facchin, & Bassetti, 2017; Pace et al., 2018). Scoring was performed by a single observer who was blinded to the group assignments of the mice. EEG epochs determined to have artefacts (interference caused by scratching, movement, eating, or drinking) were excluded from the analysis. Artefacts comprised <5-8% of all recordings used for analysis.

The numbers of epochs spent in wakefulness, NREM and REM were determined for a 24 h circadian period, across the light and dark phases (L:D = 12:12). The amount of time spent in each stage was established by the count of the epoch type (W, NR, or R) and averaged over 2-h periods. The spectral characteristics of the EEG data were further analysed. EEG power density for delta (0.5–4 Hz) frequencies in NREM sleep was computed over the 24 h of recording time. The power of each 0.5 Hz bin was first averaged across the sleep stages individually and then normalized by calculating the relative duration of each bin from the total power (0–20 Hz) for each individual animal. A two-way repeated-measures ANOVA with factors of group x time was used for the statistical analysis of two-hour averaged time-course changes in the quantity of each sleep stage (W, NREM and REM) and for physical activity. The statistical analysis of the cumulative amounts of W, NREM, and REM over the dark and light periods between genotypes was carried out with a two-tailed t-test. A two-way repeated-measures ANOVA with factors of group x frequency (Hz) was used for the statistical analysis of EEG power for NREM sleep over the 24 h recording period.

#### REM sleep characterization

Additional analyses were performed to characterize the REM sleep alterations observed in the PWScr^m+/p-^ mice. In particular, a more detailed analysis of REM sleep was carried out according to the partition of REM sleep episodes into single and sequential episodes, which has originally been proposed for rats (Zamboni, Amici, Perez, Jones, & Parmeggiani, 2001). First, the REM sleep intervals (RSIs), given by the sum of the time spent in wakefulness and in NREM sleep between two REM sleep episodes, were calculated. Here, the bimodal distribution of RSI length was determined using a kernel density estimation mathematical model, and single REM sleep episodes and sequential REM sleep episodes were discriminated (Figure S1). A single RSI is defined as one that is both preceded and followed by long RSIs >60 sec each, while sequential REM sleep episodes are those that are separated by short RSIs ≤ 60 sec each (Figure 1C).

Second, the REM-NREM cycle, which is defined as the interval between the onset of consecutive REM episodes, was assessed. The following criteria were used: 8 sec was the minimum length of a REM sleep episode; and cycles that were longer than 30 minutes were excluded from the analysis, since it has been already documented in mouse studies that the cycle length is approximately 2-5 minutes (Toth & Bhargava, 2013). An unpaired t-test was used to assess differences between short and long RSIs and differences in the number of REM-NREM sleep cycles.

#### Sleep spindle properties

Individual sleep spindles were detected using a wavelet-based spindle detection method (Bandarabadi, 2018). Briefly, the algorithm estimates the energy of EEG signal within 9-16 Hz range using the continuous wavelet transform, and then applies a threshold (3 SD + Mean) to detect candidate spindles. A lower threshold (1 SD + Mean) is then applied to find start and end points of the candidate spindles. Detected events that meet these criteria are considered as spindles; (i) duration between 0.4-2 s, (ii) number of cycles between 5-30, (ii) increase in power should be specific to the spindle range. Afterward, several properties of the detected spindles are calculated automatically, such as density, duration, central frequency, peak-to-peak amplitude, and symmetry.

#### Neurochemical analysis

Neurochemical analysis was performed in PWScr^m+/p-^ mice (n = 5) and PWScr^m+/p+^ mice (n = 5). Mice were anaesthetized with isoflurane at the beginning of the dark period to collect cerebrospinal fluid (CSF). We decided to collect samples at this time point because REM sleep alterations were observed at this stage. CSF was sampled as previously described (Liu & Duff, 2008). Mice were anaesthetized using 1.5%–2.5% isoflurane in oxygen and placed on a platform, and the arachnoid membrane covering the cisterna magna was punctured. The positive flow pressure allows the collection of CSF using a glass micropipette with a narrow tip. Three to eight microliters of CSF per mouse was collected and stored at −80 °C in polypropylene tubes.

#### CSF preparation for ultrahigh-performance liquid chromatography-tandem mass spectrometry (UPLC-MS/MS)

Eighteen microliters of a solution containing 1 µM each of Glu-d5 and GABA-d6 and 50 nM His-d4 in aqueous 0.1% formic acid was added to 2 µL of each thawed sample of CSF and then vortexed and centrifuged for 10 minutes at 14,000 rpm. The resulting supernatants were moved to glass vials for injection. Calibration standards were prepared by spiking artificial CSF (150 mM Na, 3 mM K, 1.4 mM Ca, 0.8 mM Mg, 1 mM P, 155 mM Cl, 10 mM glucose, 0.5 mg/mL albumin) with stock standard solutions. These calibrators were further diluted 1:10 with the internal standard solution, vortexed and centrifuged before injection.

#### UPLC-MS/MS

Chromatographic separation was performed with an Acquity UPLC system (Waters, Milford, MA, USA) using an ACE C18-Ar column (2 µm particle size, 2.1×150 mm (purchased from Advanced Chromatography Technologies Ltd, Aberdeen, Scotland, UK). Separation was carried out at a flow rate of 0.5 mL/min. The eluents were as follows: A, aqueous 0.1% formic acid with 5 mM n-perfluoropentanoic acid (NFPA); B, acetonitrile with 0.1% formic acid. The gradient used starts with 1 minute at 1% B, increasing linearly from 1% to 50% B in 2.5 minutes, and then remaining at 50% B for 30 sec before the column was equilibrated with the initial conditions for 2 additional minutes, adding up to a run time of 6 minutes. The column and autosampler temperatures were set to 45 and 10 °C, respectively. The injection volume was set to 4 µL. Analysis was performed on a XEVO TQ-S triple quadrupole mass spectrometer (Waters) equipped with an electrospray source operated in positive mode. Nitrogen was used as the desolvation (800 L/h, 450 °C) and collision gas. Data were acquired in the multiple reaction monitoring (MRM) mode, with the settings of the precursors, fragments, cone voltages and collision energies for each compound as indicated in Supplementary Table 1. Waters MassLynx 4.1 and TargetLynx 4.1 software were used for data acquisition and processing, respectively. A paired t-test was used to compare the levels of each neurotransmitter between the two genotypes.

### In vitro study

#### Cell culture

Cortical cultures prepared from embryonic mice at gestational day 18 were plated on 60-channel MEAs (Multi Channel Systems, MCS, Reutlingen, Germany) that had been coated with borate buffer and poly-L-lysine to promote cell adhesion (final density approximately 1200 cells/mm^2^). We recorded the activity of neuronal networks at 19 ± 1 days in vitro (DIV; mean ± SD), as already described (I. Colombi, F. Tinarelli, V. Pasquale, V. Tucci, & M. Chiappalone, 2016), in mice carrying a paternal deletion of the *Snord116* gene (PWScr^m+/p-^) (Skryabin et al., 2007) and in wild-type mice (PWScr^m+/p+^).

#### Experimental protocol

We recorded the spontaneous activity in 9 cultures from PWScr^m+/p-^ mice and 6 from PWScr^m+/p+^ mice for 2 h in culture solution as the baseline (BL) condition. Then, cells were recorded continuously for 2 hr following the administration of Carbachol (CCh, Sigma-Aldrich, 20 µM).

#### MEA recordings

Planar microelectrodes were arranged in an 8×8 layout, excluding corners and one reference electrode, for a total of 59 round planar TiN/SiN recording electrodes (30 µm diameter; 200 µm centre-to-centre inter-electrode distance). The activity of all cultures was recorded by means of the MEA60 System (MCS). The signal from each channel was sampled at 10 kHz and amplified using an MCS amplifier with a frequency band of 1 Hz-3 kHz. Each recorded channel was acquired through the data acquisition card and monitored online through MC_Rack software (MCS). In order to reduce thermal stress on the cells during the experiment, the MEAs were kept at 37 °C by means of a controlled thermostat (MCS) and covered with polydimethylsiloxane (PDMS) caps to prevent evaporation and changes in osmolality.

#### Data analysis and statistics

Starting from the raw data (i.e., the wide-band signal), we used a dual process to analyse both MUA (multiunit activity, f>300 Hz) and LFPs (local field potentials, f<300 Hz). To capture only MUA activity, we high-pass-filtered the raw signal. When spikes (i.e., single over-threshold peaks) and bursts (i.e., groups of tightly packed spikes) were detected in the MUA recordings, we computed the following parameters for each channel: mean firing rate (MFR, spikes/sec), mean bursting rate (bursts/min), inverse burst ratio (IBR, percentage of spikes outside the burst), burstiness index (BI, an index of the burstiness level of the network), burst duration (BD, msec), mean frequency intraburst (MFB, spikes/sec), interburst interval (IBI, msec) and spike time tiling coefficient (STTC). Finally, we computed the cross-correlation between each pair of burst trains recorded from active channels (i.e., with MFR>0.05 spikes/sec). For each channel, we considered only the first spike for each burst (i.e., the burst event), and we computed the cross-correlation function that represents the probability of observing a burst in one channel i at time t+T (T =3 msec) given that there is a burst in a second channel i+1 at time t. To quantify the strength of the correlation between each pair of electrodes, we evaluated the correlation peak (C_peak_). We selected only the first 100 C_peak_ values to identify only the most significant correlations. Finally, we analysed the latency from the peak (L_peak_) and considered the corresponding peak latency values of the preselected 100 strongest C_peak_ values (El Merhie et al., 2018).

To select the LFP components, we low-pass-filtered the raw data between 1 and 300 Hz. We then computed the power spectral density of the decimated signal (sampling frequency 1 kHz) (μV²/Hz) using the Welch method (windows=5 sec, overlap=50%). We considered only the lower-frequency bands of the signal; these bands, particularly the delta (0.5-4 Hz), theta (4-11 Hz), and beta (11-30 Hz) bands (I. Colombi et al., 2016), are of particular interest in the study of the sleep-wake cycle. To characterize the LFP, we calculated the power in each of those frequency bands.

### Measurement of *in vitro* and *in vivo* gene expression by real-time quantitative PCR

Total RNA was extracted from mouse cortices and from cortical cultures to assess the changes in distinct classes of genes that have already been shown to be differentially expressed during sleep and wakefulness (Cirelli, Faraguna, & Tononi, 2006; Cirelli & Tononi, 2000a, 2000b; Hinard et al., 2012). These genes are immediate early genes and clock genes (*Cfos; Arc; Homer1a, Dpb, Bmal1 and Per2*). The *Bdnf* gene, which is involved in neuroplasticity and responds rapidly to SD (Schmitt, Holsboer-Trachsler, & Eckert, 2016), was also investigated. RNA was extracted from the cortex of 10 male PWScr^m+/p-^ mice (n= 5) and PWScr^m+/p+^ mice (n= 5). Mice were sacrificed by cervical dislocation at two different time points: (i) 6 h after the onset of the light period as a baseline value (ZT6), before which the mice were kept undisturbed in their home cages and (ii) 1 h after 6 h of total SD. The latter time point was used because in the *in vitro* experiments, gene expression analysis was performed 1 h after CCh treatment. The effect of CCh resembles that of SD. Total RNA was isolated separately from the frontal cortex (FC) and parietal cortex (PC) with TRIzol (Life Technologies) according to the manufacturer’s instructions (Chomczynski & Mackey, 1995). Primary cortical neurons were treated at DIV 20 with 20 μM CCh for 1 h. Then, neurons were washed three times with ice-cold phosphate-buffered saline solution and lysed with 300 μl of TRIzol. All RNA concentrations were then determined by a NanoDrop 2000c spectrophotometer. Complementary DNA was reverse transcribed from up to 2 mg of total RNA using a high-capacity RNA-to-cDNA kit (Invitrogen) and then analysed with SYBR GREEN qPCR mix. Reactions were performed in three technical replicates using an AB 7900HT fast real-time PCR system (Applied Biosystems). The relative quantification of expression levels was performed using a previously described ΔΔCT calculation method (Pace et al., 2017). *Gapdh* was used as a reference gene. The specific primer pairs used for the analysis were designed using Primer3 (Supplementary Table 2). An unpaired t-test was used to compare differences between the genes investigated in the two genotypes.

### Statistical analysis

The normality of the distribution of values was tested with the Kolmogorov–Smirnov test. Data are presented as the mean ± standard error of the mean (SEM). As detailed above for the *in vivo* experiment, most data were analysed using two-way ANOVA and Student’s t-test as appropriate for the experimental design. If the P values reached statistical significance, Bonferroni adjustment was further applied for post hoc analysis. For the sleep analysis, Phenopy was used (Balzani, Falappa, Balci, & Tucci, 2018), while GraphPad Prism6 (GraphPad Prism Software, Inc.) was used for statistical analysis. Type I error (α was set at 0.05 (p < 0.05).

For MEA recordings, belonging to the *in vitro* experiment, data were analysed using two-way repeated-measures ANOVA (factors=genotype x treatment) followed by Tukey’s post hoc test. SigmaPlot software (Systat Software Inc.) was used for statistical analysis to compare data from two different populations, we performed the Mann-Whitney test, since data were not normally distributed.

## Results

### Paternal *Snord116* deletion impacts the microstructure of sleep in mice

To assess whether *Snord116* might contribute to sleep disturbances associated with PWS, we recorded the sleep-wake cycle over the 24 h circadian period (12 h:12 h light/dark cycle) of PWScr^m+/p-^ and PWScr^m+/p+^ mice (Figure 1A). Both genotypes exhibited the typical circadian change in the sleep-wake distribution, with reduced sleep (NREM and REM sleep) during the dark hours compared to the light hours (Figure S2A). Total sleep and wake duration showed no obvious differences between the two cohorts of mice (Figure S2A). In contrast, the total amount of REM sleep was increased in mice lacking a paternal *Snord116 gene* during the dark period (PWScr^m+/p+^ 2.425 ± 0.5083 vs PWScr^m+/p+^ 3.865 ± 0.4334, t(18)= 2.15, p=0.04; Figure 1B). Next, we assessed the EEG power spectra, and Delta (4 – 4.5 Hz) power density during NREM sleep was found to be significantly reduced in the PWScr^m+/p-^ mice compared to the control mice (two-way ANOVA: F (39, 351) = 44.13 p<0.0001; “Frequency Hz”, Figure S2B).

Since the EEG analysis highlighted an alteration of REM sleep, which reflects a change in the distribution of REM periods with no effects on total sleep or wakefulness, we further explored the duration of REM sleep episodes in detail. Our results show, for the first time, the presence of a bimodal distribution of RSI in mice. RSI was bimodally distributed in both genotypes of mice over the 24 h circadian period, with the two clusters separated at 55 sec, which accounts for the minimum frequency (Figure 1C). Based on these parameters, the two subpopulations of RSI detected were used to distinguish single REM sleep episodes (long RSI >60 sec) and sequential REM sleep episodes (short RSI ≤60 sec). Although a bimodal distribution was observed in both genotypes of mice, the frequency of short RSIs was significantly higher in the PWScr^m+/p-^ mice than in the PWScr^m+/p+^ mice (PWScr^m+/p+^ 3.40 ± 0.99 vs PWScr^m+/p+^ 7.6 ± 1.6, t(18)= 2.21, p=0.03; Figure 1C), suggesting a high REM sleep propensity in the mutant mice. Conversely, the numbers of long RSIs were unchanged between the two genotypes (Figure 1C). In addition, the REM-NREM cycle was analysed as an identifier of sleep propensity (Trachsel, Tobler, Achermann, & Borbely, 1991). Our data showed that the cycle length was approximately 2-5 minutes in both genotypes of mice, as already described (Toth & Bhargava, 2013). However, in the PWScr^m+/p-^ mice, a large increase in REM-NREM cycles was observed over the 24 h of the sleep-wake cycle (PWScr^m+/p+^ 9.00 ± 2.06 vs PWScr^m+/p+^ 15.15 ± 1.51, t(18)= 2.40, p=0.02; Figure 1C).

### Paternal deletion of *Snord116* alters sleep spindles in PWS mice

Sleep spindle properties were explored in relation to the circadian and homeostatic components of sleep (Figure 1D). Sleep spindles are the best-characterized source of EEG power in the 9–16 Hz range during NREM sleep (see Figure 1E). We calculated these spindle properties for each genotype at different time points and found that the density of spindles was unchanged between the two genotypes in all conditions investigated (Figure 1E). However, we observed that spindle duration (t(1988)= 3.04, p=0.002; Figure 1E), central frequency (t(1988)= 2.70, p=0.006; Figure 1E), and the number of spindle cycles (t(1988)= 3.47, p=0.005; Figure 1E) were significantly increased in the PWScr^m+/p-^ mutants compared to the control mice. Conversely, spindle amplitude was significantly lower in the PWScr^m+/p-^ mutants compared to the PWScr^m+/p-^ mice (t(1988)= 18.59, p<0.0001; Figure 1E).

### PWS mutants show increased spontaneous physical activity over the circadian cycle

Physical activity was assessed in awake mice without any motor constraints. During the 24 hours of the circadian period, both genotypes showed an increase in physical activity during the night and a decrease in motility during the day. Interestingly, mutant mice displayed an increase in spontaneous physical activity during the subjective night (light phase, resting phase for mice) compared to control mice (PWScr^m+/p+^ 2.48 ± 0.44 vs PWScr^m+/p+^ 3.96 ± 0.33, t(18)= 2.6, p=0.01; Figure 2A). Moreover, physical activity was also increased in the mutant mice during the first hours of the dark period compared to the motor activity of control mice (two-way ANOVA: F(11,99) = 9.318, p<0.0001, “time of day”, and F(1,9) = 5.23, p=0.04, “groups”; Figure 2A).

**Figure 2.**
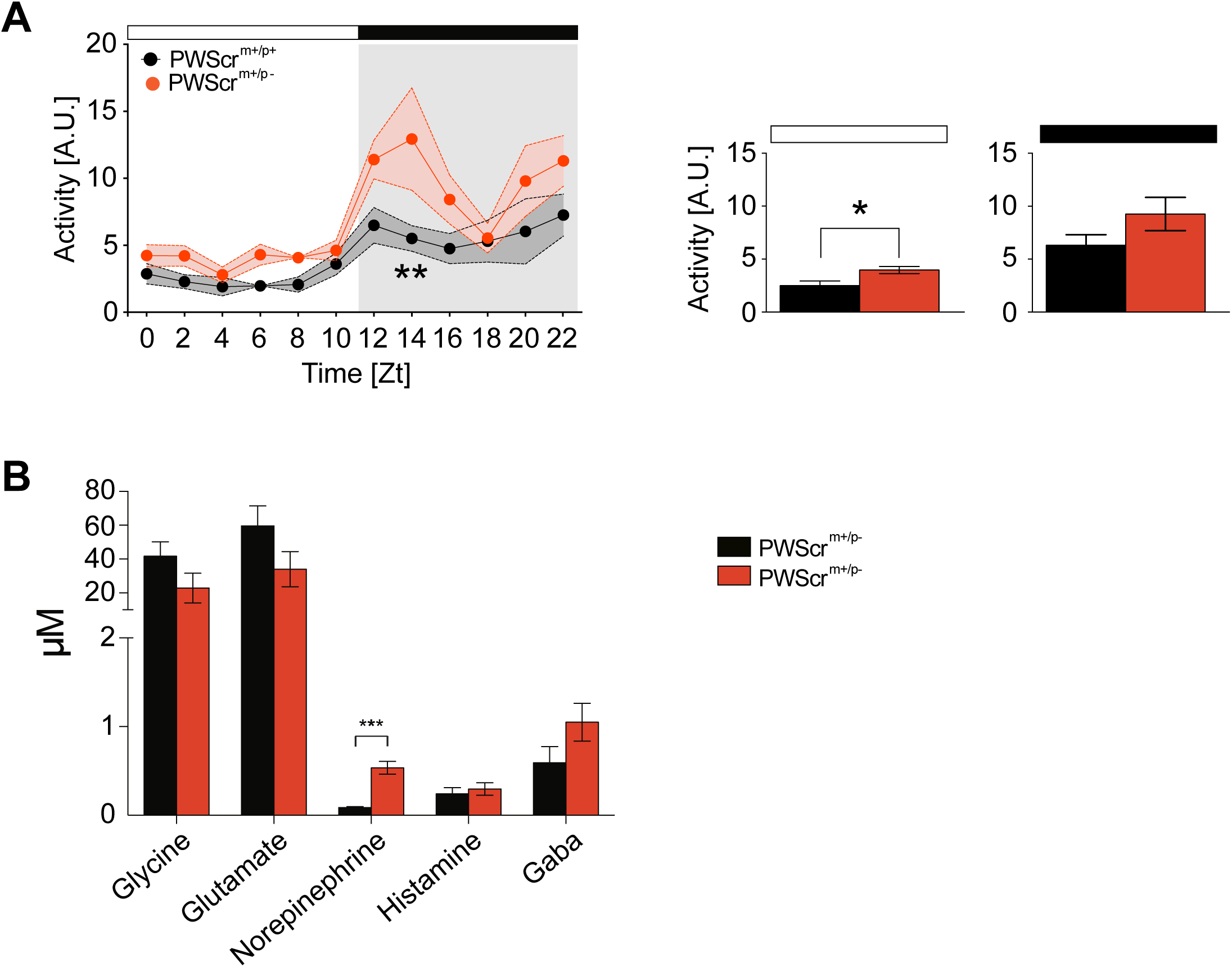
Paternal *Snord116* deletion increases physical activity and norepinephrine levels. **A)** On the left, telemetry measurements of daily activity in home cages monitored 24-hours per day in PWScrm^+/p+^ (n= 10 in black) and PWScrm^+/p-^ mice (n= 10 in red). On the right the spontaneous physical activity averaged during the 12 h light and dark period. Values are the 12 h means ± SEM. **B)** Levels of neurotransmitters (Glycine, glutamate, norepinephrine, histamine and GABA) assessed in the CSF of PWScrm^+/p+^ (n= 10 in black) and PWScrm^+/p-^ mice (n= 10 in red). Mice were sacrificed at the beginning of the dark period. Values are expressed as mean ± SEM. *p < 0.05; **p < 0.01; ***p < 0.001

### Paternal *Snord116* deletion leads to dysregulation of norepinephrine

Neuromodulators are fundamental players in the regulation of sleep-wake cycles. Here, we report an assessment of the levels of glycine, glutamate, norepinephrine, histamine and GABA in the CSF of both genotypes. Notably, among all the investigated neuropeptides, it was found that the levels of norepinephrine, a neurotransmitter that is involved in arousal and is also important for the regulation of REM sleep (Mallick et al., 2002; Mitchell & Weinshenker, 2010), was significantly increased in the PWScr^m+/p-^ mice relative to the control (PWScr^m+/p+^ 0.53 ± 0.07 vs PWScr^m+/p+^ 0.08 ± 0.01, t(8)= 6.22, p=0.0003; Figure 2B).

### REM-like *in vitro* phenotypes emerge from paternal *Snord116* deletion neurons

To assess whether *Snord116* impacts the communication of cortical networks of neurons isolated from the thalamocortical connections, we studied electrophysiological and transcriptional changes in dissociated embryonic cortical neurons obtained from PWScr^m+/p-^ and PWScr^m+/p+^ mice. We administered CCh to the cultures to suppress the delta wave activity and emphasize activity in the theta frequency range. We recorded two hours of baseline neuronal activity (i.e., basal phase) in the culture medium, followed by two hours of CCh administration for both genotypes (PWScr^m+/p-^ and PWScr^m+/p+^ cultures). We compared the electrophysiological activity on two different time scales: LFP and MUA.

First, we analysed the low-frequency component of the signal to investigate whether PWScr^m+/p-^ cultures showed an alteration in the theta oscillations during CCh administration. Our results, in agreement with our previous studies, demonstrated dysregulation of the electrophysiological activity in the theta frequency band of PWScr^m+/p-^ mice during REM sleep (Lassi et al., 2016) over the 24 h circadian period (as the baseline condition). In the baseline condition, cortical networks from both genotypes revealed spontaneous synchronized bursting resembling the slow-wave activity typical of NREM sleep. Indeed, the analysis of the LFP performed in PWScr^m+/p-^ and PWScr^m+/p+^ cultures during spontaneous phase did not indicate differences between the genotypes (Figure S3).

We then stimulated cortical cultures with CCh as previously described (I. Colombi et al., 2016) (Figure 3B) and we normalized power with respect to the basal condition. Following CCh administration, the power of the delta waves decreased by 80% in both sets of cultures (Figure 3C). We found a significant difference between the two sets of cultures (i.e., PWScr^m+/p-^ and PWScr^m+/p+^) only in the theta waves after 1 h of CCh application (time point=30′: q= 3.222; p=0.029; time point=60′ q=3.828; p=0.011; time point=90′: q= 3.94; p=0.009; time point=120′: q= 3.693; p=0.013) (Figure 3C1). Specifically, the theta power of the control cultures decreased by 60%. In contrast, the theta power of the PWS cultures decreased by 30%. These results are consistent with the results of the *in vivo* experiment that revealed abnormalities in REM sleep characterized by theta waves. Beta waves did not show significant differences between the two genotypes. Detailed statistical information about the multiple comparisons is available in Table S3.

**Figure 3.**
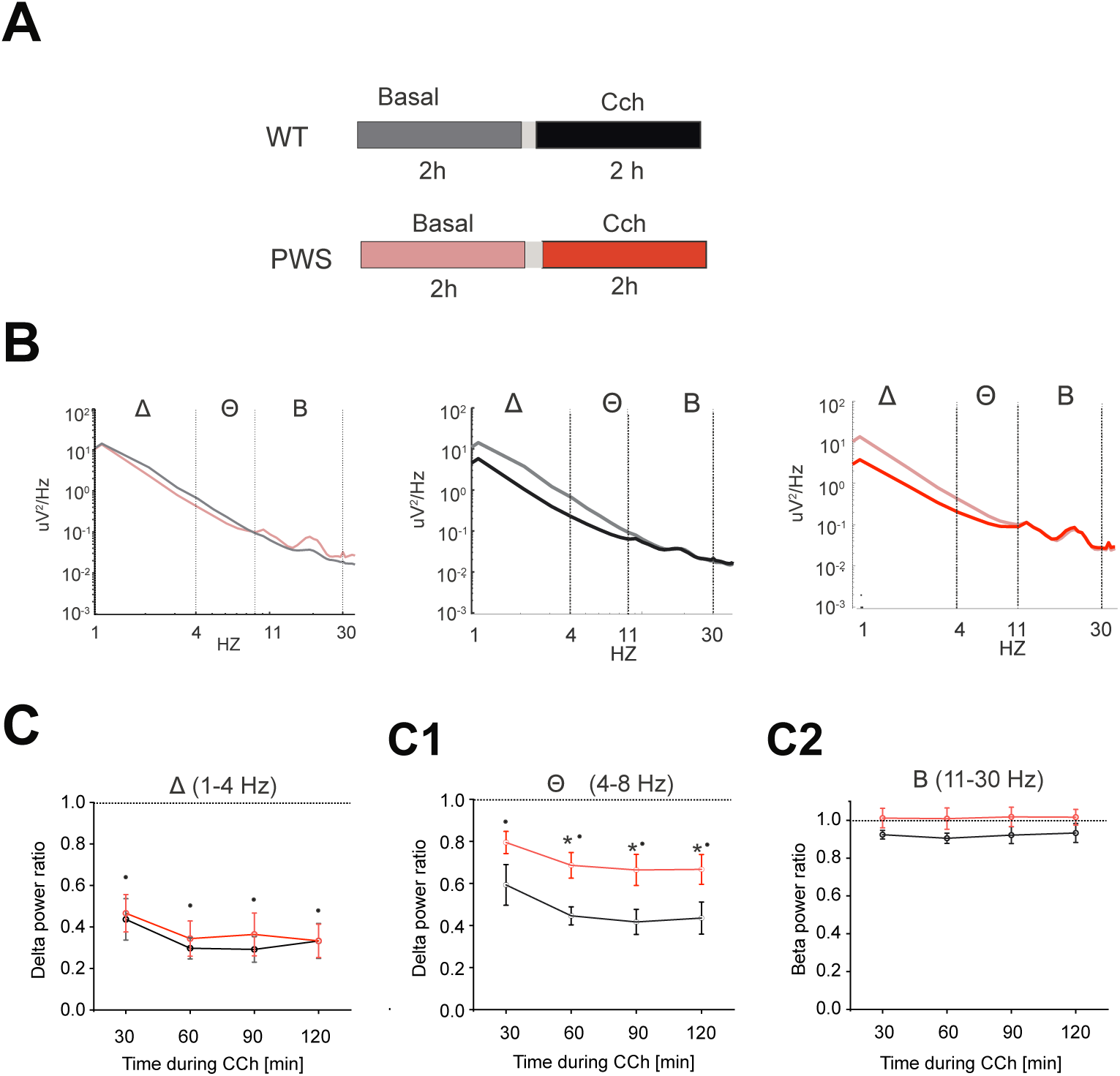
LFP analysis of PWScr ^m+/p-^ and PWScr ^m+/p+^ cultures. **A)** Experimental protocol adopted for the experiments. We recorded two hours of spontaneous phase followed by 2 hours of CCh administration. **B)** (Left) Power spectral density of basal segments for PWScr ^m+/p-^ and PWScr ^m+/p+^ cultures. We computed the power for each set of bands delta (1-4 Hz), theta (5-9 Hz) and beta (9-13 Hz). We did not find any differences during the basal phase. (Center) Power spectral density of PWScr ^m+/p+^ cultures under baseline conditions (grey line) and during CCh application (black line). CCh caused suppression of all the low waves considered. (Left panel) Power spectral density of PWScr ^m+/p-^ cultures under baseline conditions (light red line) and during CCh application (red line). **C)** Delta power following CCh administration. We normalized the power in each band with respect to the basal value (indicated by the dashed line). After dividing the entire recording into 30-minute intervals, we did not find a significant difference between PWScr ^m+/p-^ and PWScr ^m+/p+^ cultures: CCh administration caused a significantly decrease of the delta power for both genotype. **C1)** Theta power following CCh administration. We normalized the power in each band with respect to the basal value (indicated by the dashed line). After dividing the entire recording into 30-minute intervals, we found a significant difference between PWScr ^m+/p-^ and PWScr ^m+/p+^ cultures after 1 hour of treatment. The theta power of the PWScr ^m+/p+^ cultures decreased by 60%, whereas that of the PWScr ^m+/p-^ cultures decreased by 30%. **C2)** Beta power following CCh administration. We normalized the power with respect to the basal value (indicated by the dashed line). After dividing the entire recording into 30-minute intervals, we did not find a significant difference between PWScr ^m+/p+^ and PWScr ^m+/p-^. All data are presented as the mean +/-SEM. Statistical analysis was performed using Tukey’s post hoc test following two-way repeated-measures ANOVA (*p<0.05 indicated statistical difference between genotype in the different time-points; ° p <0.05 indicated statistical difference within the same genotype during Cch administration with respect to the basal condition).

### Paternal *Snord116* deletion alters the synchronization of activity and delays response to CCh treatment

Subsequently, we analysed the high frequency range of the signal to evaluate possible network activity defects correlated with the syndrome. First, we compared the first two hours of the spontaneous activity highlighted in Figure 4A. PWScr^m+/p+^ revealed highly synchronized activity during the basal phase, composed of network-wide bursts separated by periods of nearly complete quiescence, or asynchronous action potentials (Chiappalone, Vato, Berdondini, Koudelka-Hep, & Martinoia, 2007; Eytan & Marom, 2006; van Pelt, Wolters, Corner, Rutten, & Ramakers, 2004), as depicted in the raster plot of one representative experiment Figure 4B. In contrast, PWScr^m+/p-^ cultures displayed a relatively asynchronous firing pattern, as demonstrated by the following analysis. We found no significant differences by comparing the firing rate during baseline, denoting a comparable starting level of activity (Figure 4C light blue box and light red box for PWScr^m+/p+ and^ PWScr^m+/p-^ cultures, respectively). PWScr^m+/p-^ cultures displayed more desynchronized activity than PWScr^m+/p+^ resulted in a reduced BI (Mann-Whitney U test; Z=2.18, p=0.02) (BI, Figure 4C1) and STTC (Mann-Whitney test; Z=2.29, p=0.01; Figure 4C2). The analysis of the cross-correlation of burst events (Figure 4C3-C4) revealed a statistically significant difference in the latency of the top 100 connections between the two groups under basal conditions (Mann-Whitney test; Z= −1.94; p=0.049) (Figure 4C4). We did not find any significant difference in the C_peak_ values (Figure 4C3). To enable the behaviour of the cultures to be compared during CCh stimulation, we normalized the results of each experiment to the mean values obtained during basal recording. In the PWScr^m+/p+^ cultures (Figure 4F, blue line), the activity level evaluated by means of the MFR increased after CCh administration compared to the basal condition, but we did not find any significant difference. At the same time, CCh application resulted in an increased number of isolated spikes (i.e., an increased IBR value, Figure 4 F1) (two-way repeated-measures ANOVA; basal phase vs CCh 30′: q= 5.87, p=0.001; basal phase vs CCh 60′: q=6.62, p<0.001; basal phase vs CCh 90′: q=7.02, p<0.001; basal phase vs CCh 120′ q=5.639, p=0.002) and a decrease in the BI (two-way repeated-′ measures ANOVA; basal phase vs CCh 30′: q=4.338; p=0.02; basal phase vs CCh 60′: q= 6.21 p<0.001; basal phase vs CCh 90′: q= 6.603, p<0.001; basal phase vs CCh 120′: q= 6.95 p<0.001) (Figure 4F2) and STTC (two-way repeated-measures ANOVA; basal phase vs CCh 30′: q= 5.181, p=0.005; basal phase vs CCh 60′: q= 7.069, p<0.001; basal ′ phase vs CCh 90′: q= 6.666 p<0.001; basal phase vs CCh 120′: q= 6.125 p<0.001) (Figure 4F3) compared to the basal phase, indicating a loss of bursting activity and of synchronicity. Moreover, CCh causes a fragmentation of burst structures resulting from an increase in BD (Figure 4 F4) (two-way repeated measure ANOVA; basal phase vs CCh 60′: q= 5.142, p=0.006; basal phase vs CCh 90′: q= 6.02, p<0.001; basal phase vs CCh 120′: q= 6.504, p<0.001), a decrease in the peak of cross-correlation on burst events (C_peak_, Figure 4F5) (two-way repeated measure ANOVA; basal phase vs CCh 30′: q= 4.474, p=0.021; basal phase vs CCh 60′: q= 5.805, p=0.001; basal phase vs CCh 90′: 5.938 p=0.001; basal phase vs CCh 120′: q= 5.58, p=0.002) and an increase in the latency of the top 100 connections (Lat, Figure 4F6) (two-way repeated measure ANOVA; basal phase vs CCh 60′: q= 4.962, p=0.008; basal phase vs CCh 90′: 4.263 p=0.031). Indeed, the relative changes of the analysed parameters upon CCh administration in PWScr^m+/p+^ cultures are consistent with the previous results obtained in cortical cultures from embryonic rats despite the different *in vitro* ages (DIVs) of cultures recorded (I. Colombi et al., 2016). Although the starting level of spontaneous firing was comparable between genotypes (Figure 4C), CCh had a delayed effect in PWScr^m+/p-^ cultures. We found a significant difference between the first 30 minutes and the last one hour of CCh administration for the MFR ratio (two-way repeated-measures ANOVA; CCh 30′ vs CCh 120′: q= 4.783, p=0.012) (Figure 4 F). We did not find any significant difference in the IBR ratio of PWScr^m+/p-^ at any time point of 30 minutes (Figure 4F1). In contrast, the BI (two-way repeated-measures ANOVA; CCh 30′ vs CCh 90′: q= 5.37, p=0.003; CCh 30′ vs CCh 120′: q= 8.568, p<0.001) (Figure 4F2), STTCS (two-way repeated-measures ANOVA; CCh 30′ vs CCh 60′: q=5.03, p=0.007; CCh 30′ vs CCh 90′: q= 5.934, p=0.001; CCh 30′ vs CCh 120′: q= 6.501, p<0.001) (Figure 4F3), BD (two-way repeated-measures ANOVA; CCh 30′ vs CCh 90′: q= 5.544, p=0.002; CCh 30′ vs CCh 120′: q= 5.093, p=0.006) (Figure 4F4), C_peak_ Figure 4F5) (two-way repeated-measures ANOVA; CCh 30′ vs CCh 90′: q= 4.86, p=0.01; CCh 30′ vs CCh 120′: q= 5.008, p=0.007) and latency ratio (Figure 4 F6) (two-way repeated-measures ANOVA; CCh 30′ vs CCh 90′: q= 4.607, p=0.016; CCh 30′ vs CCh 120′: q= 4.244, p=0.032) showed a statistically significant difference between the time point of 30 minutes and the subsequent time points of 90 and 120 minutes by CCh treatment. Indeed, in contrast to PWScr^m+/p+^ cultures, the above parameters showed a statistically significant difference with respect to the basal conditions starting 90 minutes after the administration of the drug (Figure 4 F2-F6).

**Figure 4.**
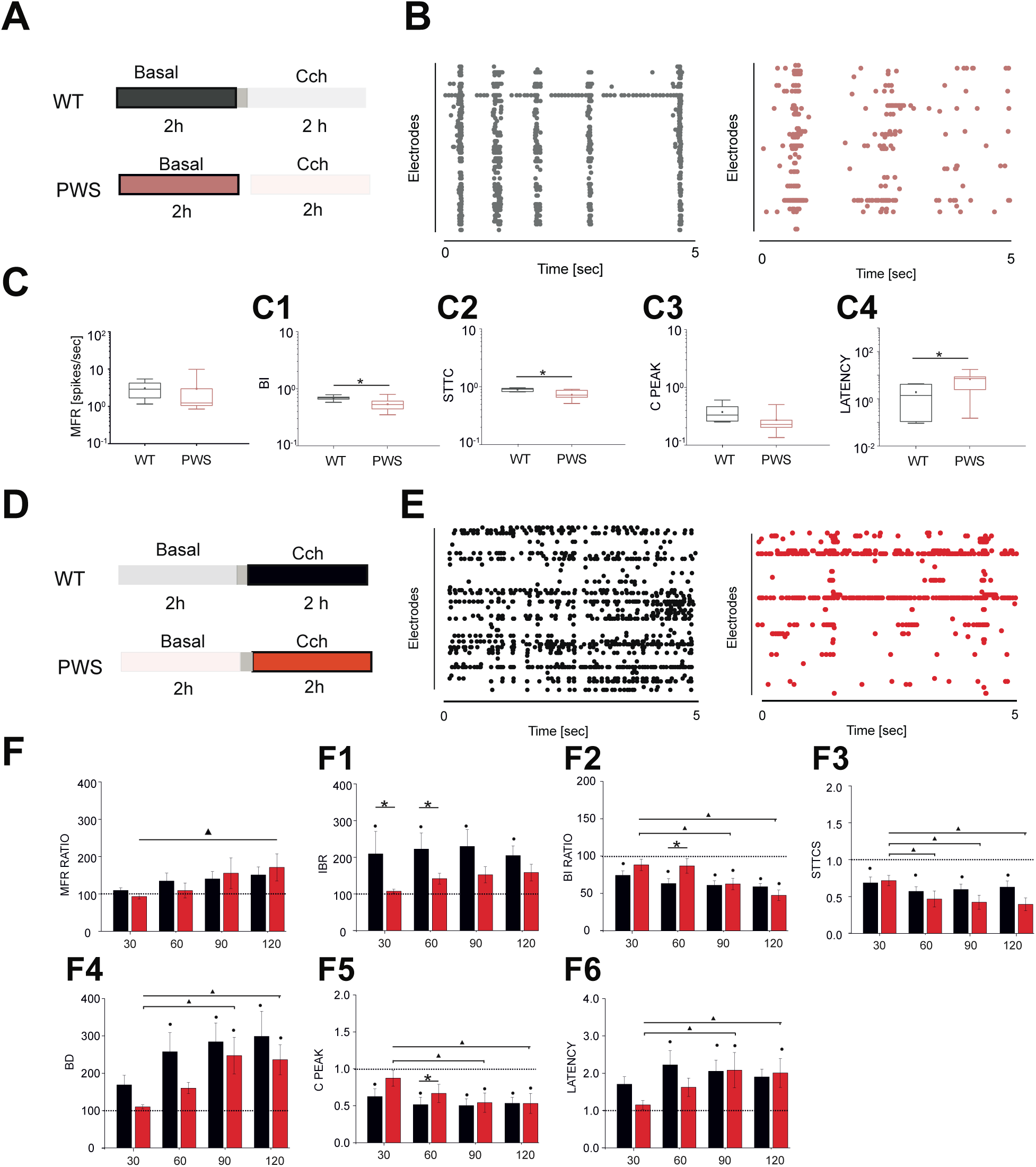
MEA analysis of PWScr ^m+/p+^ and PWScr ^m+/p-^ cortical cultures. **A)** Experimental protocol adopted for the experiments. The 2 hours of basal recording used in the analysis are highlighted. **B)** 150-s raster plots of cortical cultures at DIV 18 during basal phase for PWScr ^m+/p+^ and PWScr ^m+/p-^cultures. **C)** Comparison of MFR in PWScr ^m+/p+^ (grey box, N=6) and PWScr ^m+/p-^ (light red box, n = 9) cultures during basal phase **C1)** Comparison of BI in PWScr ^m+/p+^ (grey box plot, n = 6) and PWScr ^m+/p-^(light red box, n = 9) cultures during basal phase. **C2**) Comparison of STTC values in PWScr ^m+/p+^ (grey box plot, n = 6) and PWS (light red box, n = 9) cultures during basal phase. **C3)** Comparison of The top 100 C peak values calculated on burst events in PWScr ^m+/p+^ (grey box plot, n = 6) and PWScr ^m+/p-^ (light red box, n = 9) cultures during basal phase. **C4)** Comparison of The top 100 Latency value calculated on burst events in PWScr ^m+/p+^ (grey box plot, n = 6) and PWScr ^m+/p-^ (light red box, n = 9) cultures during basal phase. **D)** Experimental protocol adopted for the experiments. The 2 hours of CCh recording used in the analysis are highlighted. **E)** 150-s raster plots of cortical cultures at DIV 18 during basal and CCh administration for PWScr ^m+/p+^ and PWScr ^m+/p-^cultures **F)** Comparison of 30-minute time point of MFR in PWScr ^m+/p+^ (black columns, n = 6) and PWScr ^m+/p-^ (red columns, n = 9) cultures following CCh administration. **F1)** Comparison of 30-minute time point of IBR in PWScr ^m+/p+^ (black columns, n = 6) and PWScr ^m+/p-^ (red columns, n = 9) cultures following CCh administration. **F2)** Comparison of BI in PWScr ^m+/p+^ (black columns, n = 6) and PWScr ^m+/p-^ (red columns, n = 9) cultures following CCh administration. **F3)** Comparison of the top 100 STTC values obtained for PWScr ^m+/p+^ (black columns, n 6) and PWScr ^m+/p-^ (red columns, n = 9) cultures following CCh administration. **F4)** Comparison of BD in PWScr ^m+/p+^ (black column, n = 6) and PWScr ^m+/p-^ (red column, n = 9) cultures following CCh administration. **F5)** Cross-correlation analysis of burst events. Bar plot of the 100 highest Cpeak values at each 30-minute time point. **F6**) Bar plot of the corresponding peak latency values (Lpeak) of the preselected 100 highest Cpeak values. In each box plot (C C4), the small square indicates the mean, the central line indicates the median, and the box limits indicate the 25th and 75th percentiles. All data are presented as the mean +/- SEM. Statistical analysis was performed using Tukey’s post hoc test following two-way repeated-measures ANOVA (*p < 0.05 indicated statistical difference among the different time points treatment (i.e. CCh 60’ and CCh 120’) within the same genotype,° p < 0.05 indicated statistical difference within the same genotype during Cch administration with respect to the basal condition).

When we compared the behaviour of PWScr^m+/p+^ and PWScr^m+/p-^, we observed a higher increase in the IBR values for PWScr^m+/p+^ than for PWScr^m+/p-^ in the first hour by the CCh administration (Figure 5 F1) (two-way repeated measure ANOVA; CCh 30′: q= 3.767, p=0.013; CCh 60′: q= 2.986, p=0.045). At the same time, the BI ratio (Figure 4 F2) (two-way repeated-measures ANOVA; CCh 60′: q= 3.37, p=0.02) and the C_peak_ ratio (Figure 4 F5) (two-way repeated-measures ANOVA; CCh 60′: q= 3.10, p=0.03) showed lower values for PWScr^m+/p+^ than for PWScr^m+/p-^ after one hour of treatment. Instead, at the same time point, the BD ratio revealed higher values for PWScr^m+/p+^ than for PWScr^m+/p-^ (Figure 4 F4). Therefore, paternal *Snord116* deletion causes alterations in the synchronization level of activity and a delayed response to CCh treatment. All the statistical details from the multiple pairwise comparisons are available in Table S4.

**Figure 5.**
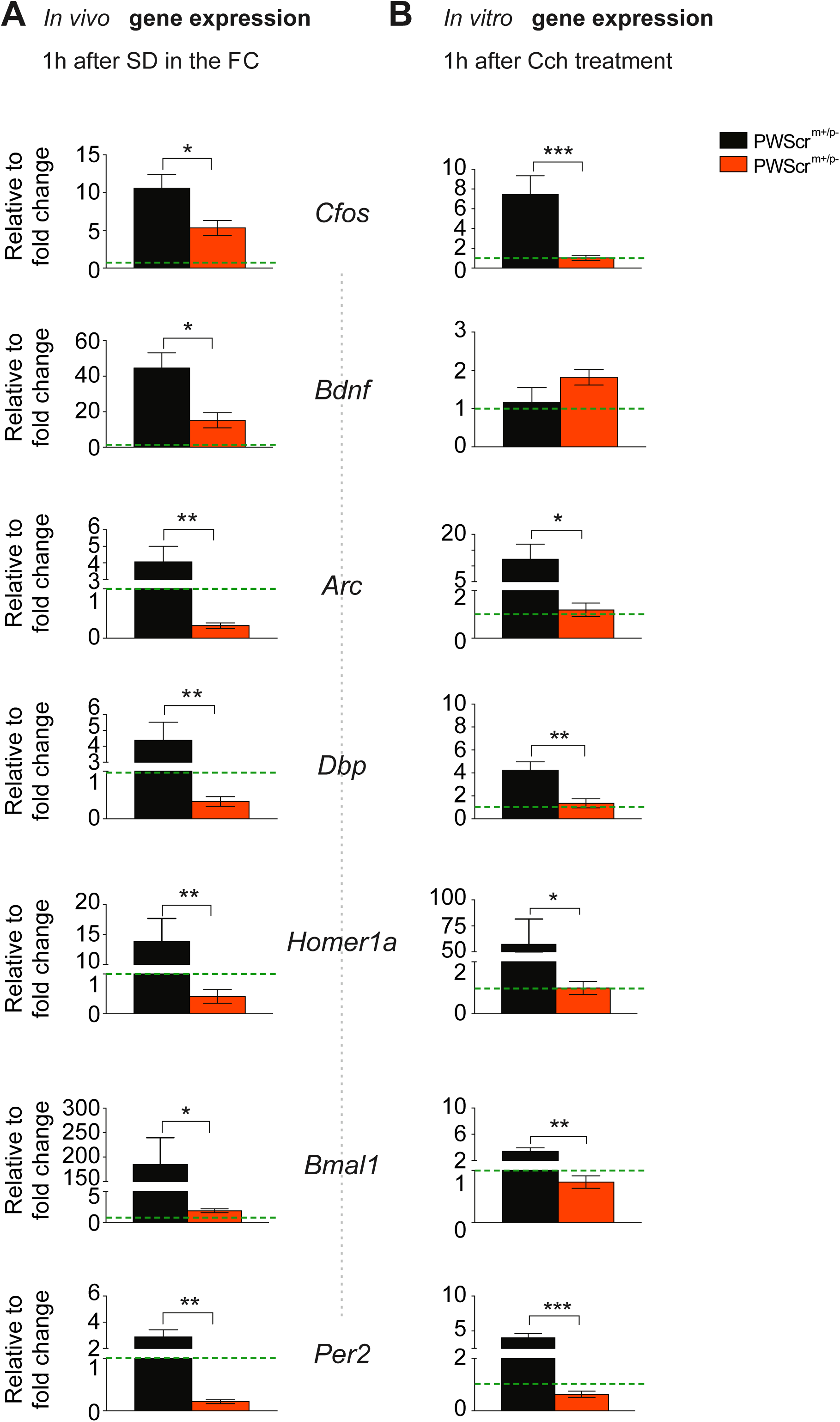
Paternal *Snord116* deletion alters the immediate early genes (IEG) in-vivo and in-vitro. **A**) IEG gene expression analysis was performed in PWScr^m+/p+^ (n= 10 in black) and PWScr^m+/p-^ mice (n= 10 in red). Mice were sacrificed after 6 hours of sleep deprivation and 1 hour of rebound. Values expressed are relative to wildtype control average ± SEM. Green line show the baseline level. **p < 0.01; ***p < 0.001. **B**) IEG gene expression analysis was performed in embryonic cortical neurons obtained from PWScr^m+/p+^ (n= 5 in black) and PWScr^m+/p-^ mice (n= 5 in red). Cells were collected after 1 hour of CCh treatment. p < 0.05; **p < 0.01; ***p < 0.001. IEG investigated: *Fos* Proto-Oncogene, AP-1 Transcription Factor Subunit (*C-fos*); Brain-derived neurotrophic factor (*Bdnf*); Activity Regulated Cytoskeleton Associated Protein (*Arc*); D-Box Binding PAR BZIP Transcription Factor (*Dbp)*; Homer Scaffold Protein 1 (*Homer1a*); Brain and Muscle ARNT-Like 1 (*Bmal1* also named as Aryl hydrocarbon receptor nuclear translocator-like protein 1 (*Arntl*)); Period Circadian Regulator 2 (*Per2*).

### Paternal *Snord116* deletion alters the expression of immediate early genes (IEGs) in vivo and in vitro

We explored the impact of *Snord116* on the expression of selected IEGs in the cortex, as they represent an important target of both neuronal responses and sleep homeostasis. In the *in vivo* experiment, IEG expression was investigated in the FC and PC of adult mice (15-18 weeks old) sacrificed at ZT6 and 1 h after 6 h of SD. We observed that at ZT6, which constitutes the baseline condition, *Bdnf* expression was significantly reduced in the PWScr^m+/p-^ mice compared to the control (PWScr^m+/p+^ 1 ± 0.12 vs PWScr^m+/p+^ 0.46 ± 0.07, t(8)= 3.73, p=0.005; Figure S4A) in the PC. This increase in the *Bdnf* level was also observed in the PC of mutant mice (PWScr^m+/p+^ 1 ± 0.12 vs PWScr^m+/p+^ 0.62 ± 0.06, t(8)= 2.68, p=0.02; Figure S4A). Moreover, in the PC, a significant reduction in *Cfos* (PWScr^m+/p+^ 1 ± 0.37 vs PWScr^m+/p+^ 0.09 ± 0.03, t(8)= 2.38, p=0.04; Figure S4A) and *Arc* (PWScr^m+/p+^ 1 ± 0.19 vs PWScr^m+/p+^ 0.37 ± 0.13, t(8)= 2.64, p=0.02; Figure S4A) genes was also observed. We noticed that, 1 h after 6 h of SD, the expression levels of all investigated IEGs were significantly reduced in the mutant mice compared to control mice (Figure 5; see Table S2 for statistical analysis). Conversely, the expression levels of all investigated IEGs were, as expected, significantly increased compared to the baseline value (ZT6, non-SD mice) in the PWScr^m+/p+^ mice (Hinard et al., 2012) (Figure 5; see Table S3 for statistical analysis). The same scenario was observed for the expression of IEGs in the PC of the mutant mice (Figure S4A; see Table S3 for statistical analysis).

Subsequently, the alteration in cortical IEG expression observed in living PWScr^m+/p-^ mice 1 h after SD was investigated to determine whether it is also reflected in the embryonic cortical neurons. For the *in vitro* analyses, cortical cultures were stimulated with 20 µM of CCh, and RNA was extracted 1 h later, which resembled the *in vivo* SD condition. Conversely, the ZT6 treatment in the *in vivo* experiment mirrors the cell culture stimulated with water (baseline). In primary cell cultures at baseline, the only IEG whose expression was significantly different between the two genotypes was *Per2.* In this condition, *Per2* was found to be significantly higher in the PWScr^m+/p-^ cultures than in the control cultures (PWScr^m+/p+^ 1 ± 0.31 vs PWScr^m+/p+^ 19± 2.6, t(8)= 7.01, p=0.0001; Figure S4B). It was also observed that, 1 h after CCh treatment, cells displayed the same alterations in IEG expression that were observed in the *in vivo* experiment. Specifically, it was observed that all IEG expression levels were significantly lower in the PWScr^m+/p-^ mice than in the control mice (Figure 5; see Table S4 for statistical analysis). In contrast, PWScr^m+/p+^ mice showed significantly increased IEG expression after CCh treatments, as observed in the *in vitro* experiments (Figure 5; see Table S4 for statistical analysis). Interestingly, only *Bdnf*, which is not an IEG but is involved in the plasticity mechanisms, did not undergo a change in expression in cultured cortical neurons 1 h after CCh (Figure 5; see Table S4 for statistical analysis), while in living mice, the expression of this gene was significantly altered before and after SD.

Overall, these *in vivo* and *in vitro* results show that IEG expression in the cortex is significantly affected by the loss of the *Snord116* gene. Additionally, it was observed herein that the wake-like (CCh treatment vs SD mice) and sleep-like states (water treatment vs ZT6 mice) *in vitro* underwent changes in gene expression that resembled those in the cerebral cortex of living animals.

## Discussion

The results of this study confirm that the *Snord116* gene significantly affects REM sleep occurrence and its regulation. Sleep spindle properties were found to be altered in the PWScr^m+/p-^ mice, while their numbers were unchanged between the two genotypes, suggesting dysfunctions in the cortex where sleep spindles are amplified and not in the thalamus where they are generated. Moreover, we also described a previously unreported form of intrinsic cortex dysfunction observed in primary cortical neurons obtained from PWScr^m+/p-^ mice, reinforcing our results obtained on sleep spindles. Next, the electrophysiological and molecular changes that we observed in primary cortical neurons resemble the same sleep phenotypes and transcriptional changes observed in living mice.

Mice carrying a paternal deletion of the *Snord116* gene display REM sleep dysregulation. The increase in REM sleep is pronounced during the dark period, when mice are most active, and may resemble the EDS observed in humans PWS (Camfferman et al., 2008). Since it has already been reported that REM sleep is precisely regulated in terms of its duration, while NREM is substantially regulated in terms of intensity (Amici et al., 2008; Borbely & Achermann, 1999), we decided to analyse the succession of REM sleep episodes and the period of time intervening between the end of one REM sleep episode and the beginning of the next (REM sleep interval, RSI). RSI follows a bimodal distribution characterized by short and long periods. The presence of a bimodal distribution has already been described (Amici, Zamboni, Perez, Jones, & Parmeggiani, 1998) in many species, such as rats (Amici et al., 1994), cats (Ursin, 1970) and humans (Merica & Gaillard, 1991), but, at the best of our knowledge, no data are yet available on the RSI distribution in mice. For the first time, our data reveal a bimodal RSI distribution in mice, with the minimum frequency at 55 sec, while in rats, the minimum between the two peaks has been identified at 3 min (Amici et al., 1998). A bimodal distribution of RSI was observed in both genotypes of mice. However, mutant mice showed a significant increase in short RSI. Short RSI has been described as an indicator of the capacity to produce REM sleep in accordance to the homeostatic drive under favourable ambient conditions and associated with the capacity to produce REM sleep proportionally to the homeostatic drive under favourable ambient conditions (Amici et al., 1998; Zamboni, Perez, Amici, Jones, & Parmeggiani, 1999). Indeed, the amount of REM sleep with short RSIs has been shown: i) to be increased during a REM sleep rebound in proportion to the degree of previous REM deprivation induced by low ambient temperature (Zamboni et al., 2001); ii) to be depressed when animals are kept under an uncomfortable ambient condition, such as during the exposure to a low Ta (Amici et al., 1998), and concomitantly with a reduced capacity of cAMP accumulation at hypothalamic-preoptic level (Jones et al., 2008; Zamboni et al., 2004). This observation may reflect a strengthened drive for REM sleep and/or a distorted and more favourable perception of the ambient conditions, in terms of the thermal comfort, in the mutant mice. Therefore, since sleep propensity also affects the NREM-REM cycle, we also assessed the number of these cycles over the 24 h of the BL in both genotypes of mice. As expected, mutant mice presented a significant increase in the NREM-REM cycle (approximately 2-5 minutes) (Toth & Bhargava, 2013) compared to the control group, suggesting, once again, that the mutant mice have a strong tendency to fall asleep easily. Overall, these data imply that the loss of *Snord116* significantly compromised sleep microstructure, mainly affecting REM sleep.

We found that PWScr^m+/p-^ mice showed alterations in REM sleep without affecting NREM sleep. An exception, however, was found in the sleep spindles, hallmarks of NREM sleep that reflect the activity of complementary thalamocortical circuits. The thalamic reticular nucleus (TRN) is the spindle pacemaker, and TRN/thalamus circuits can generate spindles in isolation, although cortical inputs may contribute by initiating or amplifying spindle oscillations (Ferrarelli & Tononi, 2017; Fuentealba & Steriade, 2005). Here, we show that the duration, amplitude and frequency of sleep spindles are significantly altered in PWScr^m+/p-^ mice, while their numbers are unchanged between the two genotypes. This data suggests that TRN/thalamus circuits or in corticothalamic afferents to these intra-thalamic circuits may probably not affected in the mutant mice, while, an impairment of cortical amplification may explain the alterations of sleep spindles properties. Since it has been shown that sleep spindles facilitate neuroplasticity and support learning, memory consolidation, and intellectual performance (Gruber & Wise, 2016), and since sleep spindle alterations have been documented in children with neurodevelopmental disorders (Burnett et al., 2017; Gruber & Wise, 2016), we speculate that the neurodevelopmental and cognitive alterations observed in these mice (Adhikari et al., 2018) may also arise from altered thalamocortical input. Thus, we believe that sleep spindle alterations may either reflect the severity of the underlying disorder or directly exacerbate the severity of impairments. Evidence also indicates that alterations in sleep spindle properties have a very early association with an increased risk of cognitive impairment (Taillard et al., 2019); our results imply that sleep spindles may also represent reliable sleep EEG biomarkers associated with PWS disorder, although a better characterization of these findings in PWS subjects is needed.

Additionally, we observed that PWScr^m+/p-^ mice have dysregulated norepinephrine levels in the CSF collected at the beginning of the dark period. Norepinephrine is a neurotransmitter important for maintaining normal sleep states, and dysregulated norepinephrine signalling is responsible for cataplexy attacks, which, together with narcolepsy, represent common features of PWS (Weselake et al., 2014). Thus, the activity of norepinephrine neurons in the locus coeruleus is important in modulating cortical activity during NREM sleep (Eschenko, Magri, Panzeri, & Sara, 2012). Norepinephrine not only influences general arousal but also affects locomotor activity. Indeed, it has been observed that intraventricular infusion of norepinephrine increases motor activity (Geyer, Segal, & Mandell, 1972). Interestingly, we observed that norepinephrine was significantly higher in the mutant mice than in the controls, suggesting that norepinephrine may also have a role in controlling motor activity in these mice. Overall, these data imply that mutant mice may also have a dysregulated norepinephrine system. The extent to which norepinephrine alterations may affect the sleep-wake cycle and locomotor activity should be further investigated and may have therapeutic relevance.

Our data imply a dysfunction of cortical amplification, manifested by the altered sleep spindles properties not affecting their numbers observed in the mutant mice. Based on these results, we investigated whether cortical neurons isolated from thalamocortical projections showed an altered intrinsic mechanism of sleep comparable to the sleep endophenotypes observed in living mice. To address this question, we used an *in vitro* model. Previous studies have demonstrated that embryonic cortical neurons on MEA are able to recapitulate some essential features of sleep in a controllable way (I. Colombi et al., 2016; Hinard et al., 2012; Saberi-Moghadam, Simi, Setareh, Mikhail, & Tafti, 2018). Here, we treated cortical cultures with CCh to change the synchronized default sleep-like state, characterized by slow-wave oscillations typical of NREM sleep, into a theta-predominant state, which is typical of REM sleep. Because of this EEG feature of REM, the shorthand definition of REM sleep is a highly activated brain in a paralysed body (O’Malley & Datta, 2013). The LFP analysis of MEA cortical neurons revealed abnormalities in the theta waves in the PWScr^m+/p-^ neurons similar to those observed in living mice. CCh administration caused evident desynchronization of the activity of both genotypes, but mutant cultures showed different response profiles after CCh treatment compared to control cultures. In particular, PWScr^m+/p-^ cultures displayed less reactivity in response to the treatment, especially during the first hour. Overall, these results suggested that PWS cultures displayed alterations in neuronal activity patterns during spontaneous activity and when external stimulation was applied. Additionally, the analysis of the MEA frequency range revealed aberrant neuronal network activity patterns. In particular, PWScr^m+/p-^ cultures displayed less synchronized activity than control cultures did, as indicated by their lower BI and STTCs. Moreover, the analysis of the cross-correlation during the basal period showed shorter latencies for PWScr^m+/p+^ cultures than for PWScr^m+/p-^ cultures. Taken together, these data imply that *Snord116* may either directly influence the neuronal synchronization of cortical neurons or affect it indirectly via norepinephrine, which we found to be altered in our study in living mice. Indeed, norepinephrine is a neurotransmitter that decreases network synchrony both *in vivo* (Bergles, Doze, Madison, & Smith, 1996; Colonnese et al., 2010) and *in vitro* (Bergles et al., 1996), consistent with our results, in which PWScr^m+/p-^ cultures showed an increased level of this neurotransmitter that may be responsible for the increased desynchronization level of activity in the MEA recording.

Finally, we also assessed the expression of IEGs in both *in vitro* and *in vivo* models, and we observed that our *in vitro* model exactly recapitulates the transcriptional alteration observed in living mice. This finding also suggests that cortical cultures coupled to MEA represent a promising tool to identify novel therapeutic targets, such as sleep feature alterations and network synchronization defects. Moreover, these data suggest, in agreement with a recent publication (Coulson et al., 2018), that the *Snord116* gene affects the transcriptional profiles of circadian genes in the cortex (*Per2* and *Bmal1*), which are involved in development outside the suprachiasmatic nucleus and contribute to brain plasticity (Kobayashi, Ye, & Hensch, 2015). Indeed, we also found that the PWScr^m+/p-^ mice had low levels of *Bdnf* mRNA, which is extremely important in the regulation of synapses and the plasticity process. This latter evidence may pave the way for new interventional approaches for PWS by using TrkB agonists or by using compounds that increase the BDNF level (Habtemariam, 2018).

Overall, our results suggest that *Snord116* is important in controlling REM sleep. Additionally, here we provide the first evidence supporting the role of *Snord116* in regulating cortical neuronal activity, opening avenues for new interventions in PWS. The dysregulation of sleep spindles in our mutant mice raises the possibility that this phenomenon can be a clinically useful marker to assess PWS symptoms in humans.

## Author contributions

MP and VT designed the study. MP, MF performed the animal experiments, assisted by AF. IC performed the MEA experiment on cells. VT and MC provided infrastructural support. AF and MP performed the gene expression analysis. MP, MF, AF, IC, analyzed the data. MB supervised by AA performed sleep spindles analysis. MP, IC, MC, MC, RA and VT drafted and finalized the manuscript. All authors revised and finalized it.

## Acknowledgement

We thank Kenda Alau, a master student of Tucci’s laboratory for her help with the RNA extraction and RT-qPCR. AA and MEBA received funding from the European Union’s Horizon 2020 Research and Innovation Programme under Grant Agreements No. 696656:Graphene Flagship, Core 1 and No. 785219:Graphene Flagship, Core 2

## Notes

Conflict of Interest: each author discloses the absence of any conflicts of interest relative to the research covered in the submitted manuscript.

