## Supplementary figures and images for "The paternally imprinted gene *Snord116* regulates cortical neuronal activity"

### Figure S1

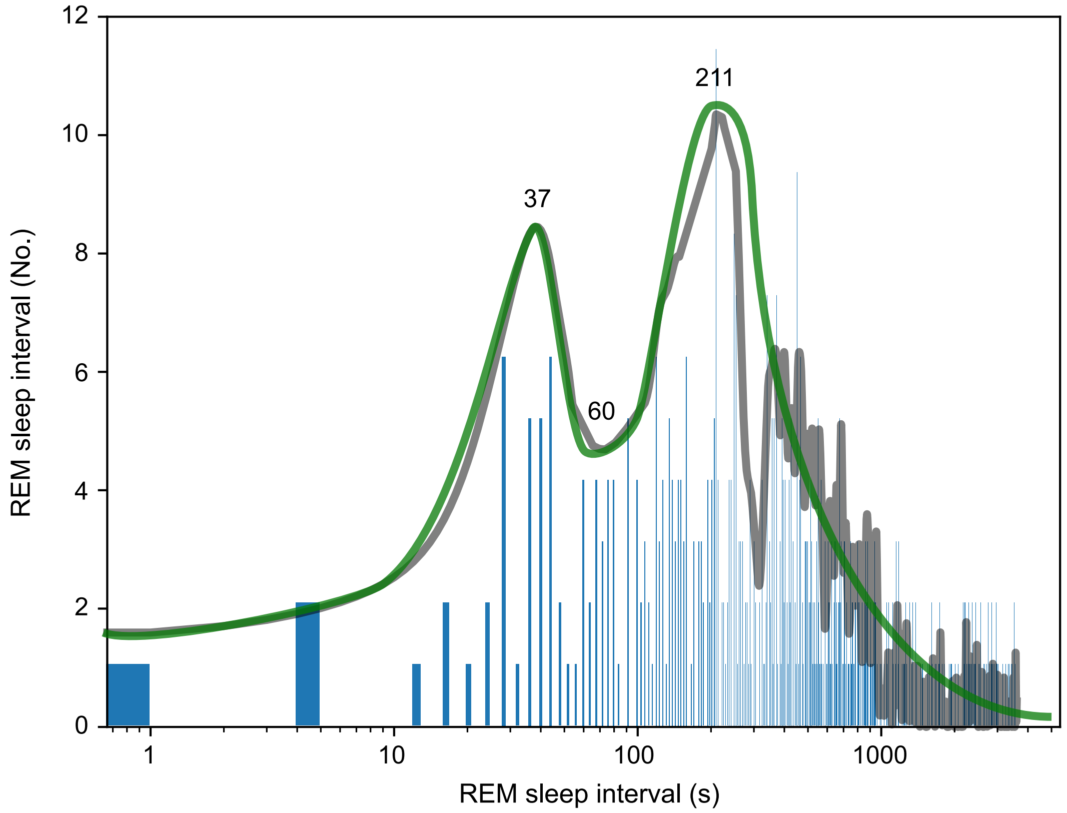

### Figure S2

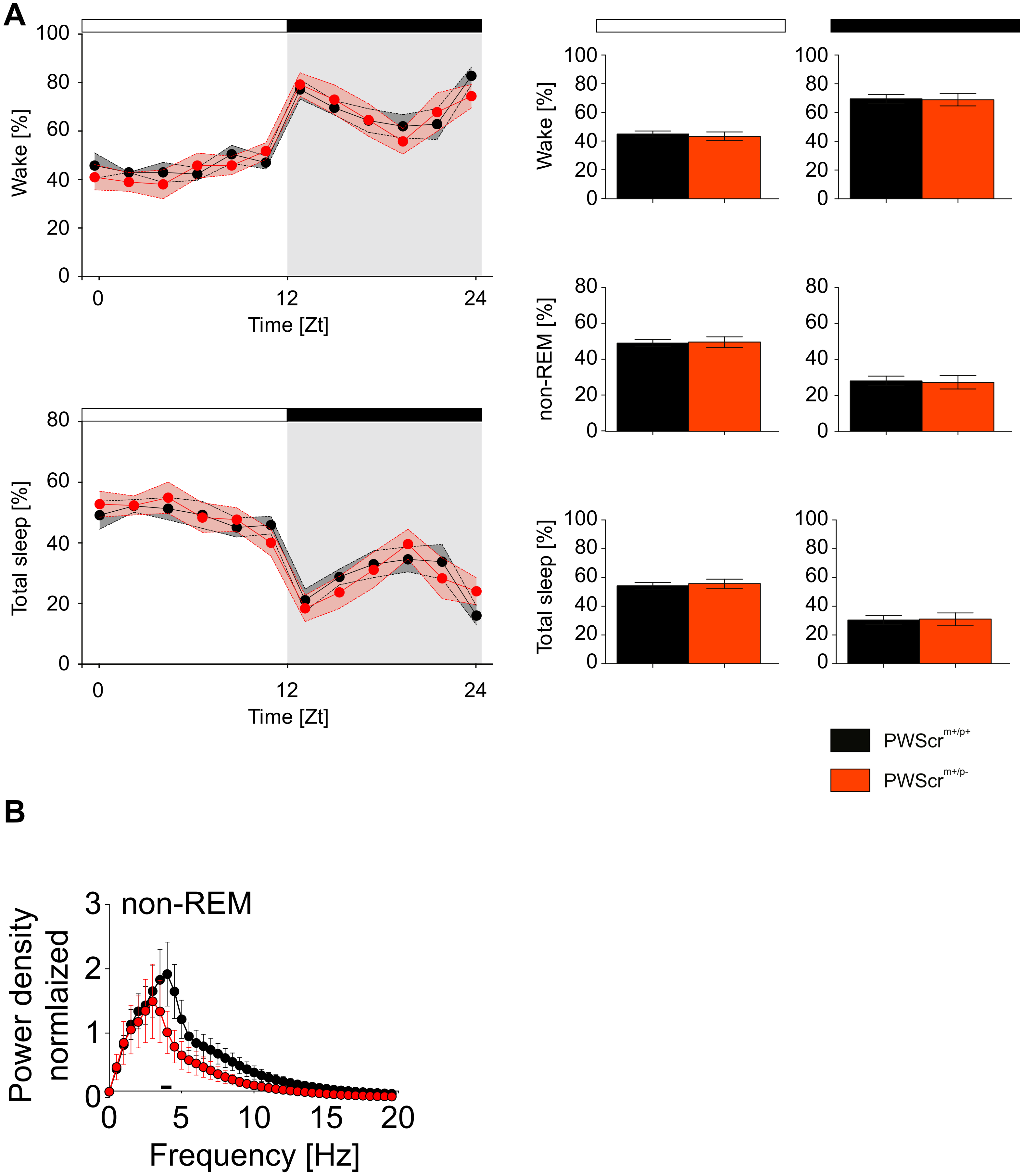

### Figure S3

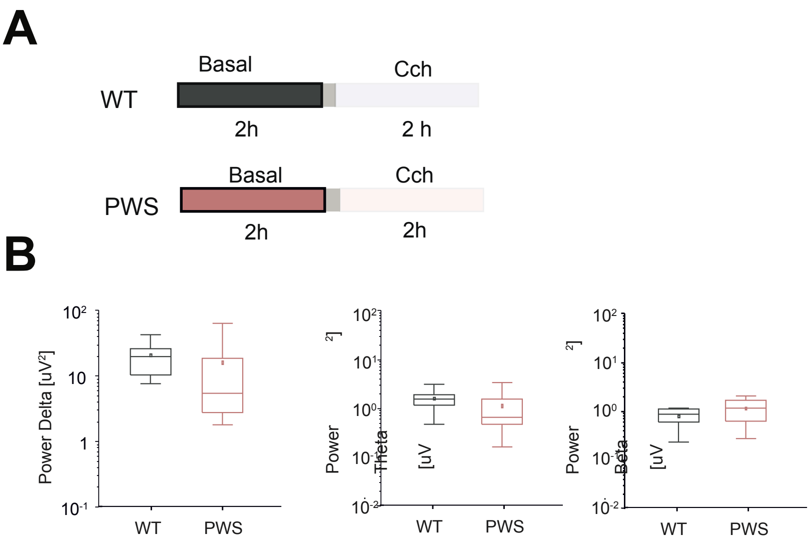

### Figure S4

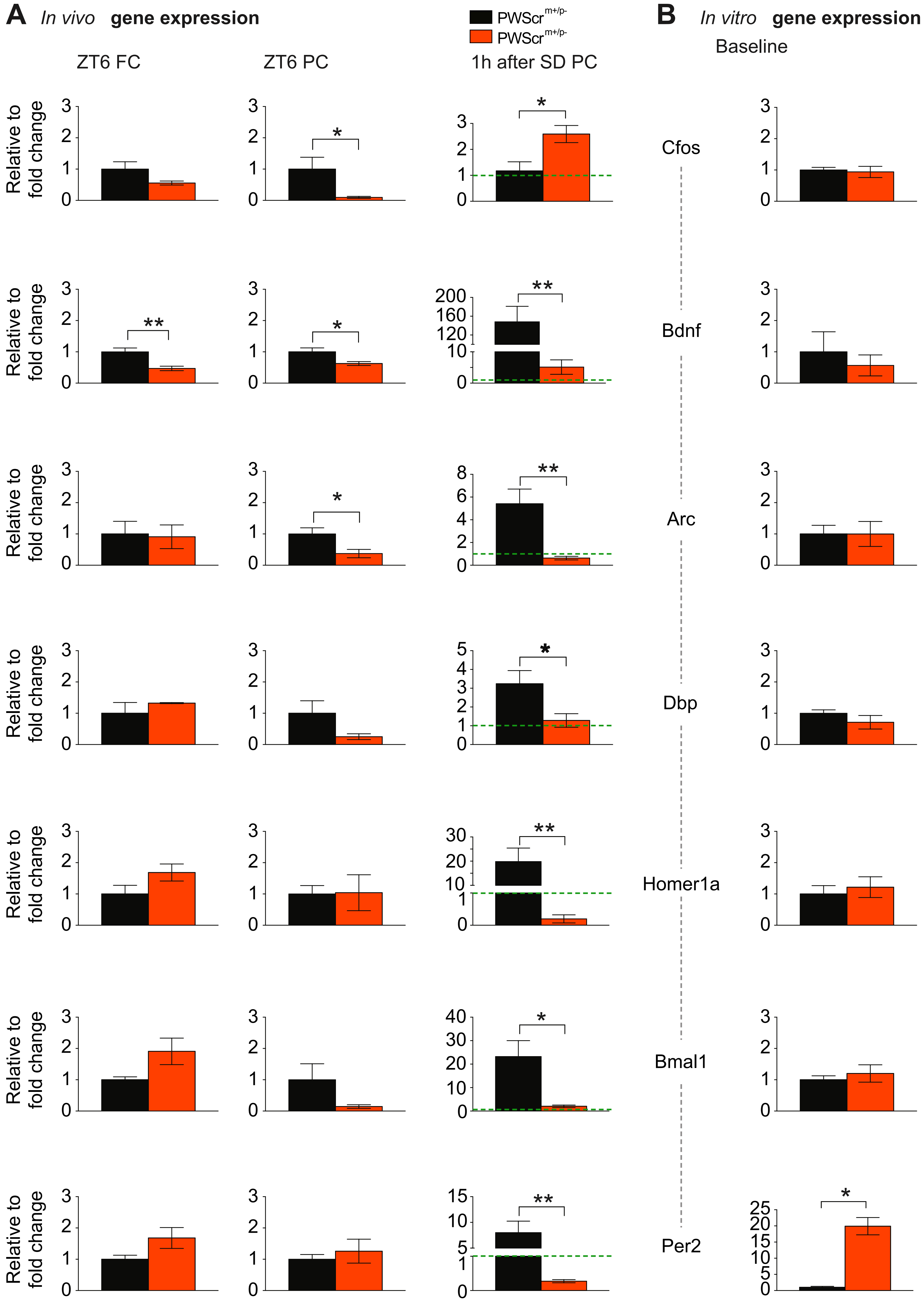
