## Supplementary materials for "The paternally imprinted gene *Snord116* regulates cortical neuronal activity"

**Conflict of Interest: each author discloses the absence of any conflicts of interest relative to the research covered in the submitted manuscript.**

**Methods**

Genotyping

To determine their genotypes, ear punches were performed at the age of 4 weeks for DNA detection using PCR. Deletion of PWScr resulted in a PCR product of 300 bp, which was absent in the WT genotype. To genotype mice, PCR analysis of genomic DNA from ear punches were performed using the primer pair PWScrF1/ PWScrR2 (5’-AGAATCGCTTGAACCCAGGA and 5’-GAGAAGCCCTGTAACATGTCA, respectively). PCR cycling conditions were as follows: 94°C – 2 min, 35 cycles of 94°C – 30 sec, 55°C – 30 sec, 72°C – 30 sec, followed by a final extension of 72°C – 9 min

**Table S1. MRM settings for the neurotransmitters and isotopically labeled internal standards. Transitions used for quantification are indicated in bold.**

| Matrix | Analyte | MRM Transition | Cone voltage (V) | Collision energy (eV) |
| --- | --- | --- | --- | --- |
| CSF | GLY | **76 > 30** | **15** | **5** |
|  |  | 76 > 48 | 15 | 5 |
|  | Glu | **148 > 84** | **15** | **15** |
|  |  | 148 > 130 | 15 | 10 |
|  | Glu d5 | 153 > 99 | 20 | 20 |
|  | GABA | 104 > 69 | 20 | 15 |
|  |  | **104 > 87** | **20** | **10** |
|  | GABA d6 | 110 > 93 | 20 | 10 |
|  | NE | **152 > 107** | **25** | **15** |
|  |  | 152 > 135 | 25 | 10 |
|  | EP | **166 > 107** | **30** | **20** |
|  |  | 166 > 135 | 30 | 15 |
|  | DA | 154 > 91 | 15 | 20 |
|  |  | **154 > 137** | **15** | **10** |
|  | ACh | 87 > 43 | 40 | 35 |
|  |  | **146 > 87** | **20** | **15** |
|  | 5HT | 177 > 115 | 15 | 25 |
|  |  | **177 > 160** | **15** | **10** |
|  | His | 112 > 68 | 20 | 20 |
|  |  | **112 > 95** | **20** | **10** |
|  | His d4 | 116 > 99 | 20 | 15 |
|  | MHis | **126 > 68** | **20** | **15** |
|  |  | 126 > 109 | 20 | 5 |
| Brain | His | **112 > 95** | 20 | 15 |
|  | MHis | **126 > 97** | 20 | 15 |

**Table S2 list of primers for gene expression analysis**

| Gene | Primer | Primer Sequence |
| --- | --- | --- |
| c-Fos | Fos_qRT F  Fos_qRT R | 5’ – GGGGACAGCCTTTCCTACTA – 3’  5’ – CTGTCACCGTGGGGATAAAG – 3’ |
| Bdnf | Bdnf_qRT F  Bdnf_qRT R | 5’ – ACACTGAGTCTCCAGGACAGCAA – 3’  5’ – AAATAACCATAGTAAGGAAAAGGATGGT – 3’ |
| Arc | Arc_qRT F  Arc_qRT R | 5’ – CCCTGCAGCCCAAGTTCAAG – 3’  5’ – GAAGGCTCAGCTGCCTGCTC – 3’ |
| Dbp | Dbp_qRT F  Dbp_qRT R | 5’ – CGTGGCGGTGCTAATGACCTT – 3’  5’ – CATGGCCTGGCCTGCTTGA – 3’ |
| Homer1A | Homer1A_qRT F  Homer1A_qRT R | 5’ – GCATTGCCATTTCCACATAGG – 3’  5’ – ATGAACTTCCATATTTATCCACCCTTACTT – 3’ |
| Bmal1 | Bmal1_qRT F  Bmal1_qRT R | 5’ – CCGTGCTAAGGATGGCTGTT – 3’  5’ – TTGGCTTGTAGTTTGCTTCTGTGT – 3’ |
| Per2 | Per2_qRT F  Per2_qRT R | 5’ – AGCTACACCACCCCTTACAAGCT – 3’  5’ – GACACGGCAGAAAAAAGATTTCTC – 3’ |
| Gapdh | Gapdh_M_qRT F  Gapdh_M_qRT R | 5’ – GAACATCATCCCTGCATCCA – 3’  5’ – CCAGTGAGCTTCCCGTTCA – 3’ |

**Table S3. The significance of immediate early gene (IEG) expression in-vivo experiment**

|  | PWScrm^+/p+^mice  Vs  PWScrm^+/p−^mice  Baseline | PWScrm^+/p+^mice  Vs  PWScinrm^+/p−^mice  1h after CCh |
| --- | --- | --- |
| *C-fos* | *n.s* | t(8)= 2,31 p= .04 |
| *Bdnf* | *n.s* | *n.s* |
| *Arc* | *n.s* | t(8)= 2,31 p= .04 |
| *Dbp* | *n.s* | t(8)= 3.50 p= .008 |
| *Homer1a* | *n.s* | t(8)= 2,30 p= .04 |
| *Bmal1* | *n.s* | t(8)= 4,71 p= .001 |
| *Per2* | t(8)= 7,01 p= .0001 | t(8)= 5,50 p= .0006 |

**Table S4. The significance of immediate early gene (IEG) expression in-vitro experiment**

| Genes | PWScrm^+/p+^mice  Vs  PWScrm^+/p−^mice  ZT6 FC | PWScrm^+/p+^mice  Vs  PWScinrm^+/p−^mice  ZT6 PC | PWScrm^+/p+^mice  Vs  PWScrm^+/p−^mice  After 6h of SD FC | PWScrm^+/p+^mice  Vs  PWScrm^+/p−^mice  After 6h of SD FC |
| --- | --- | --- | --- | --- |
| *C-fos* | *n.s* | t(8)= 2,38 p= .04 | t(8)= 2,53 p= .03 | t(8)= 2,95 p= .01 |
| *Bdnf* | t(8)= 3,73 p= .005 | t(8)= 2,68 p= .02 | t(8)= 3.97 p= .01 | t(8)= 4,32 p= .002 |
| *Arc* | *n.s* | t(8)= 2,64 p= .02 | t(8)= 3.99 p= .004 | t(8)= 3.67 p= .006 |
| *Dbp* | *n.s* | *n.s* | t(8)= 3.48 p= .008 | t(8)= 2.48 p= .03 |
| *Homer1a* | *n.s* | *n.s* | t(8)= 3.45 p= .008 | t(8)= 3.49 p= .008 |
| *Bmal1* | *n.s* | *n.s* | t(8)= 3.35 p= .01 | t(8)= 3.11 p= .01 |
| *Per2* | *n.s* | *n.s* | t(8)= 4,89 p= .001 | t(8)= 3,44 p= .008 |

**Table S5: Statistical analysis on MEAs recordings during basal recording**

| Parameter | Test |  |
| --- | --- | --- |
| STTCs | Mann-Whitney | U=47; Z=2.2981; p=0.01758 |
| BI | Mann-Whitney | U=46; Z=2.18025; p=0.02557 |
| Latency | Mann-Whitney | U=10; Z=-1.94454; p=0.04955 |

**Table S6: Statistical analysis on MEAs recordings during CCh administration**

| Parameter | Statistical test | Genotype | Treatment | Genotype X Treatment |
| --- | --- | --- | --- | --- |
| MFR | 2-Way repeated measure ANOVA | ns | F=4,452; p=0.004 | ns |
| IBR | 2-Way repeated measure ANOVA | F= 4,795;p= 0.049 | F= 9,119; p=<0.001 | F= 3,325; P= 0.018 |
| STTCs | 2-Way repeated measure ANOVA | ns | F= 25,933;p= <0.001 | ns |
| BI | 2-Way repeated measure ANOVA | ns | F=20,245; p= <0.001 | F= 2,734; p= 0.040 |
| BD | 2-Way repeated measure ANOVA | ns | F= 12.73; p= <0.001 | ns |
| C peak | 2-Way repeated measure ANOVA | ns | F= 12,510;p= <0.001 | ns |
| Latency | 2-Way repeated measure ANOVA | ns | F= 8,451, p <0.001 | ns |
| Power Theta | 2-Way repeated measure ANOVA | F= 7,418; P= 0.017 | F= 5,699; P= 0.002 | ns |

**Legends**

**Figure S1**

Whit blue bar are reported the frequency distribution of the duration of the interval from the end to one REM sleep episodes to the beginning of the subsequent episodes (REM sleep interval, RSI) during light dark cycle in mice. The grey line is the kernel density estimation constructed with the distribution of the REM sleep episodes. The green line is the smoothing of the grey line using the Savitzky-Golay Filter.

**Figure S2**

Sleep wake distribution in mice having the paternal delition of the *Snord116* **A)** On the left the time-course changes of wakefulness and total sleep in PWScrm^+/p−^ mice (in red) versus controls (in black), recorded over an uninterrupted 24 h period in 12 h light/dark cycle. Data are shown as percentage of each 2 h block spent in REM sleep within each mouse, averaged within genotypes (mean ± SEM; n= 10 PWScrm^+/p+^ and n= 10 PWScrm^+/p-^ mice). On the right the percentage of time spent in wakefulness, NREM sleep and total sleep during the 12 h light and dark period. Values are the 12 h means ± SEM. **B)** Power density during NREM sleep in PWScrm^+/p−^ mice (in red) versus controls (in black), recorded over an uninterrupted 24 h circadian period in 12 h light/dark cycle. The average EEG spectra were normalized to total EEG power from 1–20 Hz in 0.5-Hz bins. Values are expressed as mean ± SEM. *p < 0.05; **p < 0.01; ***p < 0.001

**Figure S3**

LFP analysis of PWS and WT cultures. A) Experimental protocol adopted for the experiments. The 2 hours of basal recording used in the analysis are highlighted. B) Power spectral density of basal segments for WT and PWS cultures. We computed the power for each set of bands delta (1-4 Hz), theta (5-9 Hz) and beta (9-13 Hz).We did not found any significant difference between genotypes.

**Figure S4**

Paternal Snord116 deletion alters the immediate early genes (IEG) in-vivo and in-vitro. **A**) The first two rows, IEG gene expression analysis was assessed in the frontal cortex (FC) and parietal cortex (PC) of PWScrm^+/p+^ (n= 10 in black) and PWScrm^+/p−^mice (n= 10 in red). Mice were sacrificed after 6 hours from the lights-on (ZT6). The third row, IEG gene expression analysis assessed in the parietal cortex (PC) of PWScrm^+/p+^ (n= 10 in black) and PWScrm^+/p−^mice (n= 10 in red). Mice were sacrificed after 6 hours of sleep deprivation and 1 hour of rebound. Values expressed are relative to wildtype control average ± SEM. Green line show the baseline level. **B**) IEG gene expression analysis was performed in embryonic cortical neurons obtained from PWScrm+/p+ (n= 5 in black) and PWScrm+/p- mice (n= 5 in red; p < 0.05; **p < 0.01; ***p < 0.001). IEG investigated: Fos Proto-Oncogene, AP-1 Transcription Factor Subunit (*C-fos*); Brain-derived neurotrophic factor (*Bdnf*); Activity Regulated Cytoskeleton Associated Protein (*Arc*); D-Box Binding PAR BZIP Transcription Factor (*Dbp*); Homer Scaffold Protein 1 (*Homer1a*); Brain and Muscle ARNT-Like 1 (Bmal1 also named as Aryl hydrocarbon receptor nuclear translocator-like protein 1 (*Arntl*)); Period Circadian Regulator 2 (*Per2*).
